# Mating Induces Switch From Hormone-Dependent to – Independent Steroid Receptor-Mediated Growth in *Drosophila* Prostate-Like Cells

**DOI:** 10.1101/533976

**Authors:** Aaron Leiblich, Josephine E. E. U. Hellberg, Aashika Sekar, Carina Gandy, Siamak Redhai, Mark Wainwright, Pauline Marie, Deborah C. I. Goberdhan, Freddie C. Hamdy, Clive Wilson

## Abstract

Male reproductive glands like the mammalian prostate and the paired *Drosophila melanogaster* accessory glands secrete seminal fluid components that enhance fecundity. In humans, the prostate grows throughout adult life, stimulated by environmentally regulated endocrine and local androgens. We previously showed that in each fly accessory gland, secondary cells (SCs) and their nuclei also grow in adults, a process enhanced by mating and controlled by bone morphogenetic protein (BMP) signalling. Here we demonstrate that BMP-mediated SC growth is dependent on the receptor for the developmental steroid, ecdysone, whose concentration reflects socio-sexual experience in adults. BMP signalling regulates ecdysone receptor (EcR) levels post-transcriptionally, partly via EcR’s N-terminus. Nuclear growth in virgin males is ecdysone-dependent. However, mating activates genome endoreplication to drive additional BMP-mediated nuclear growth via a cell type-specific form of hormone-independent EcR signalling. In virgin males with low ecdysone levels, this mechanism ensures resources are conserved. However, by switching to hormone-independence after mating, this control is overridden to hyper-activate growth of secretory secondary cells. Our data suggest parallels between this physiological, behaviour-induced switch and altered pathological signalling associated with prostate cancer progression.

## Introduction

In all higher organisms where fertilisation takes place in the female reproductive tract, males not only deliver sperm to females, but also transfer seminal fluid, containing a cocktail of molecules that optimise fecundity. For example, secretions from the mammalian prostate and seminal vesicles contribute most of the seminal fluid volume, activate sperm [1] and promote embryo implantation [2]. The paired accessory glands (AGs) in the fruit fly, *Drosophila melanogaster*, perform related functions and can also substantially alter female behaviour after mating, increasing egg laying, promoting sperm storage and reducing female receptivity to subsequent mating attempts [3–6]. Most of the key accessory gland proteins (Acps) involved, such as Sex Peptide, which plays a central role in driving female post-mating responses, are secreted by about 1000 so-called main cells (MCs) found in the mono-layered AG epithelium [7,8]. However, secondary cells (SCs), a small population of about 40 epithelial cells at the distal tip of each AG (Fig. 1A), also play an essential role [9–11].

As humans age, the prostate epithelium frequently becomes hyperplastic. Indeed, many males over the age of 65 develop symptomatic benign prostatic hyperplasia [12]. SCs are not proliferative, but they also grow in adults, unlike other cells in the AG [9]. Mating enhances this growth, which is most easily assayed by measuring nuclear size. Interestingly, we have found that autocrine BMP signalling is crucial for the normal, age-dependent growth of SCs both in virgin and mated males [9]. Growth of SCs involves elevated synthesis of macromolecules including secreted proteins. Inhibition of BMP signalling specifically in adult SCs reduces the ability of males to suppress female re-mating. Furthermore, BMP signalling promotes secretion of the contents of large dense-core granule-containing compartments [13] and of exosomes, nano-vesicles formed inside endosomal compartments that are released by fusion of these compartments to the plasma membrane [14]. These exosomes appear to be involved in female behavioural reprogramming, providing at least part of the explanation for SCs’ BMP-dependent effects on females.

Although BMP signalling is implicated in mammalian prostate development [15], and cancer growth and metastasis [16,17], steroid signalling through the androgen receptor (AR) is thought to be the central regulator of these processes and prostate hyperplasia [18,19]. Aberrations in steroid signalling are implicated in both benign and malignant disease of this organ [20,21]. Since androgen levels are modulated by factors such as nutrition [22] and sexual activity [23], this endocrine input potentially allows males to adapt prostate function during development, and in response to the environment and reproductive demands. In advanced cancer, hormone deprivation therapy effectively blocks tumour growth, but typically within two years, hormone-independent cells emerge, which frequently still require the AR for growth [24].

Flies employ a more limited range of steroid hormones than mammals with the major characterised steroid hormone, 20-hydroxyecdysone (usually called 20HE or ecdysone), primarily involved in developmental transitions, particularly during metamorphosis [25,26]. However, ecdysone levels also fluctuate in adult males in response to socio-sexual interactions [27]. Ecdysone regulates male courtship behaviour [28–30], and affects the male germ line [31,32]. Ecdysteroids can also induce expression of multiple Acps in AGs [33], and the ecdysone receptor (EcR) is involved in AG development [34]. However, the cells and molecular mechanisms involved in these processes, as well as the physiological functions of the EcR in adult AGs, remain unclear. We hypothesised that ecdysone signalling might affect SC function, providing a socio-sexual environmental input, which complements mating-dependent growth of SCs.

Here we show that the ecdysone receptor (EcR) is specifically expressed in SCs within the adult AG epithelium, and that EcR signalling in these cells is critical for normal growth. BMP signalling promotes growth by regulating EcR levels post-transcriptionally. While nuclear growth in virgin males is BMP-, EcR- and ecdysone-dependent, most of the EcR-mediated nuclear growth observed after mating is hormone-independent and specifically drives endoreplication. This novel form of steroid receptor control in flies permits mating and socio-sexual experience cues to be flexibly co-ordinated in regulating SC activity, hence employing resources according to demand.

## Results

### The EcR-B1 isoform is expressed exclusively by SCs in the adult AG epithelium

In order to mark and genetically manipulate SCs, we used the SC-specific esg^ts^F/O GAL4 system [35]. It can be activated exclusively in adults through inactivation of a ubiquitously expressed, temperature-sensitive form of the GAL4 inhibitor GAL80 (tub-GAL80^ts^) by a temperature shift to 28.5°C at eclosion [9,35]. Staining AGs with a pan-EcR antibody that cross-reacts with all three characterised EcR isoforms [36] revealed EcR expression in the nuclei of muscle cells, and SCs, which like MCs, are binucleate (Fig. 1B, C). This staining was lost in SCs expressing a previously characterised RNAi targeting transcripts for all EcR isoforms (Fig. 1D) [37].

**Figure 1.**
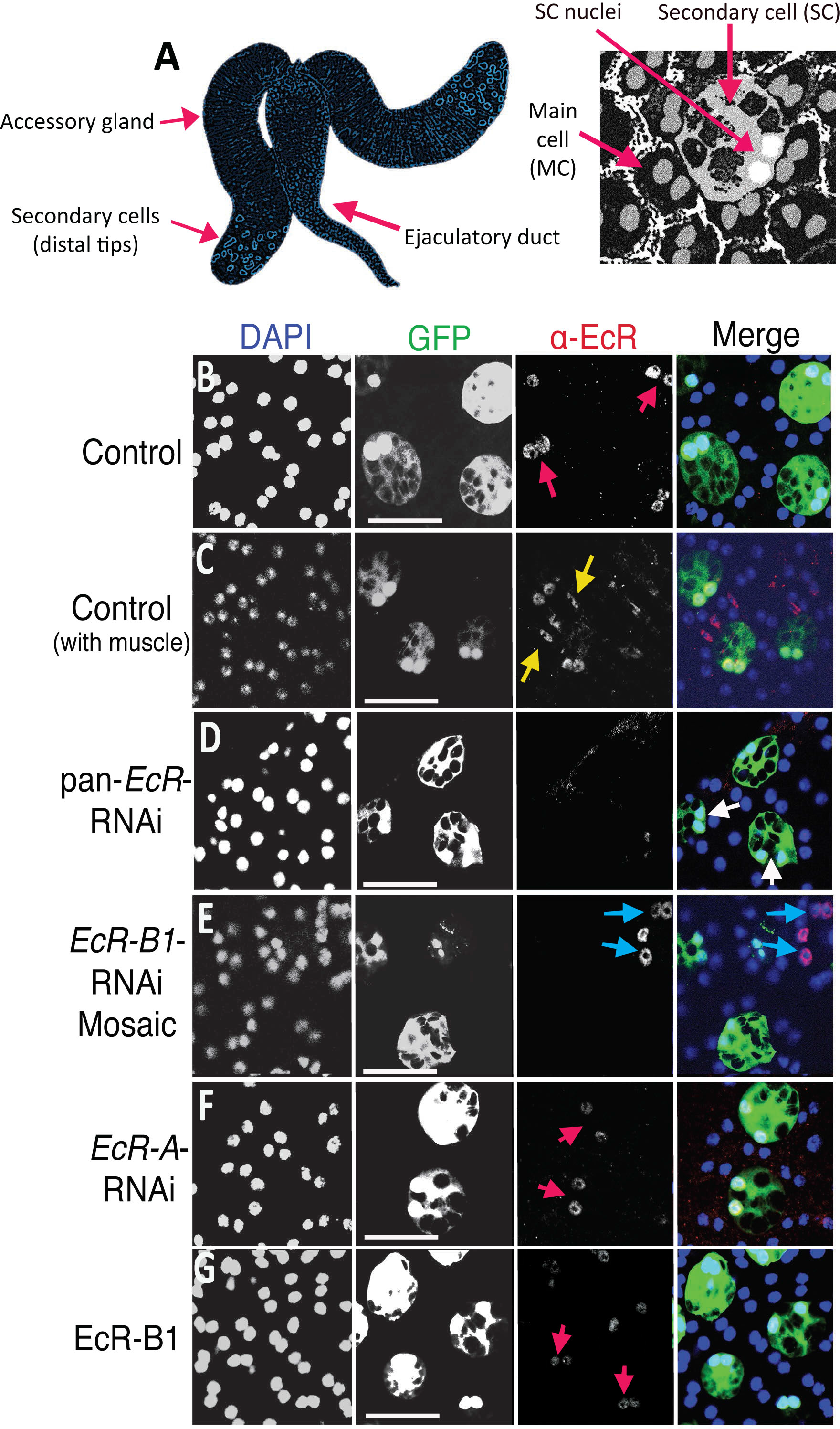
The EcR-B1 isoform of the Ecdysone Receptor is expressed in SC nuclei. A. Schematic of *Drosophila* male accessory glands and binucleate secondary and main cells within their monolayer epithelium. B-H. Images show distal tips of AGs dissected from 6-day-old males (except for mosaic in D). SCs (nuclei marked by red arrows) express nuclear GFP (which also labels SC cytosol) and other transgenes under esg^ts^F/O control. Nuclei are stained with DAPI (blue). B, C. Immunostaining with an antibody that cross-reacts with all EcR isoforms reveals expression in SC nuclei (B; red arrows) and in muscle cell nuclei (C; yellow arrows), but not in main cells (non-GFP-positive nuclei in B) in 6-day-old males. D-F. While expression of an RNAi targeting the EcR-A transcript does not affect EcR expression (F), mosaic expression of an RNAi targeting *EcR-B1* transcripts (E, green cells) or expression of an RNAi targeting all isoforms (D) obliterates nuclear EcR staining in SCs. Staining is still present in non-RNAi-expressing SCs, which are not labelled with GFP (E, blue arrows). G. esg^ts^F/O–driven expression of the EcR-B1 isoform has no detectable effect on nuclear EcR levels in SCs. Scale bars, 50 μm.

We were unable to detect a robust signal in the AG with isoform-specific EcR-A and EcR-B1 antibodies. However, in an alternative approach, we expressed isoform-specific RNAi constructs [38] in SCs under esg^ts^F/O control, either throughout adulthood, or by using a temperature-shift in 3-day-old males. The latter approach leads to maintained RNAi expression in only a subset of SCs [9]. *EcR-A*-RNAi did not affect EcR levels (Fig. 1F), but little, if any, EcR protein was detected in SCs expressing an *EcR-B1*-RNAi construct, demonstrating that EcR-B1 is the major isoform produced by these cells (Fig. 1E).

### The EcR promotes SC nuclear growth via hormone-dependent and ‒independent mechanisms

To test the roles of the EcR in adult SC growth, the pan-*EcR*-RNAi construct was expressed in these cells post-eclosion. As described previously, we assessed growth by measuring SC nuclear size in adult virgin males after 6 days relative to nuclear size in adjacent MCs, which do not grow with age or in response to mating, since this controls for nuclear size changes produced by flattening AGs upon mounting [9].

The growth of SC nuclei was inhibited by expression of *EcR*-RNAi (Fig 2A, B and F), but not by expression of a control RNAi targeting the *ry* gene (Fig. S1H). Steroid receptors typically modulate gene expression in the presence of their ligands, but in some contexts, it has been proposed that the unliganded EcR is repressive, and this repression is released by hormone [39]. To test whether ecdysone is required to induce EcR-dependent growth in SCs, we employed a temperature-sensitive allele of *ecdysoneless*, *ecd*^*1*^ [25], a gene with pleiotropic effects on processes required for ecdysone synthesis and signalling in flies [40]. When males were shifted to the non-permissive temperature (28.5°C) directly after eclosion, SCs and their nuclei failed to grow (Fig. 2E and F), mirroring the phenotype of *EcR* knockdown. This supports the hypothesis that ecdysone is required for normal EcR-dependent SC growth in virgin males.

**Figure 2.**
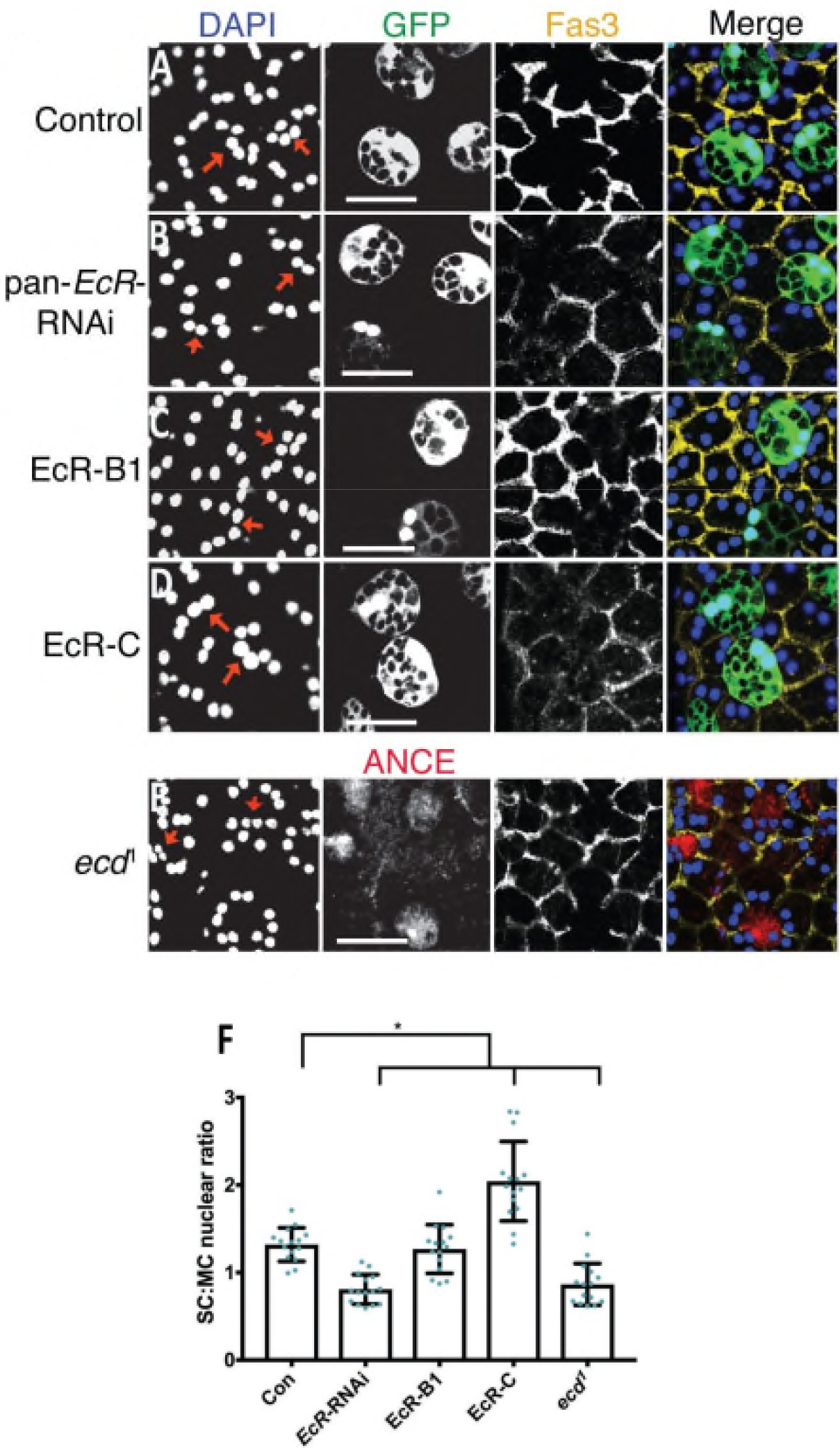
Ecdysone and the EcR are required to promote SC nuclear growth in virgin males. A-E. Dissected accessory glands from 6-day-old virgin males were stained with an antibody against Fasciclin3 (Fas3) to mark the apical outlines of SCs and neighbouring MCs (yellow) and with DAPI (blue nuclei). Selected SC nuclei are marked with red arrows and in A-D express GFP and other transgenes under esg^ts^F/O control. A, B. RNAi knockdown of EcR expression in SCs with a pan-EcR-RNAi significantly restricts growth of SC nuclei (B) compared to control glands expressing GFP only (A). C. Overexpression of EcR-B1 in SCs has no effect on SC nuclear size. D. EcR-C overexpression promotes SC nuclear growth. E. The SC nuclei of the temperature-sensitive *ecd*^*1*^ mutant are significantly smaller than control glands when adult virgin males are maintained at 28.5°C, which blocks *ecd* function. Mutant glands were co-stained with an antibody against the SC-specific secreted protein ANCE. F. Histogram showing size of SC nuclei relative to MC nuclei in *ecd*^*1*^ mutant males and in AGs where SCs express different transgenes. Significance was assessed by two-way ANOVA. *P<0.01, n>10. Scale bars, 50 μm.

In several other cell types, the EcR functions as a heterodimer with the nuclear receptor Ultraspiracle (Usp) [41,42]. Usp can promote nuclear localisation of EcR in Chinese hamster ovary cells [43]. Staining with an antibody which recognises Usp [44] revealed that this protein is expressed in SC nuclei (Fig. S2H). However, *Usp*-RNAi knockdown in SCs had no effect on cell growth (Fig. S1 A, B and H) or EcR localisation and expression (Fig. S2A, B) in SCs, even though it strongly reduced Usp levels (Fig. S2I), suggesting that conventional EcR/Usp-mediated transcriptional regulation does not drive SC growth.

To test whether EcR and ecdysone signalling is also required for the additional growth of SC nuclei observed in multiply-mated males [9], we cultured individual newly eclosed males with 7-10 virgin females for six days, then analysed nuclear size. *EcR* knockdown strongly suppressed nuclear growth under these conditions, mirroring the effects of blocking BMP signalling by expressing the transcriptional repressor Dad (Fig. 3A) [13]. However, surprisingly, temperature-shifted *ecd*^*1*^ males exhibited higher levels of SC growth after mating than mated controls (Fig. 3A-C). This suggests that unlike in virgin males, EcR-mediated growth in response to mating is primarily hormone-independent, and indeed, ecdysone blocks signalling under these conditions.

**Figure 3.**
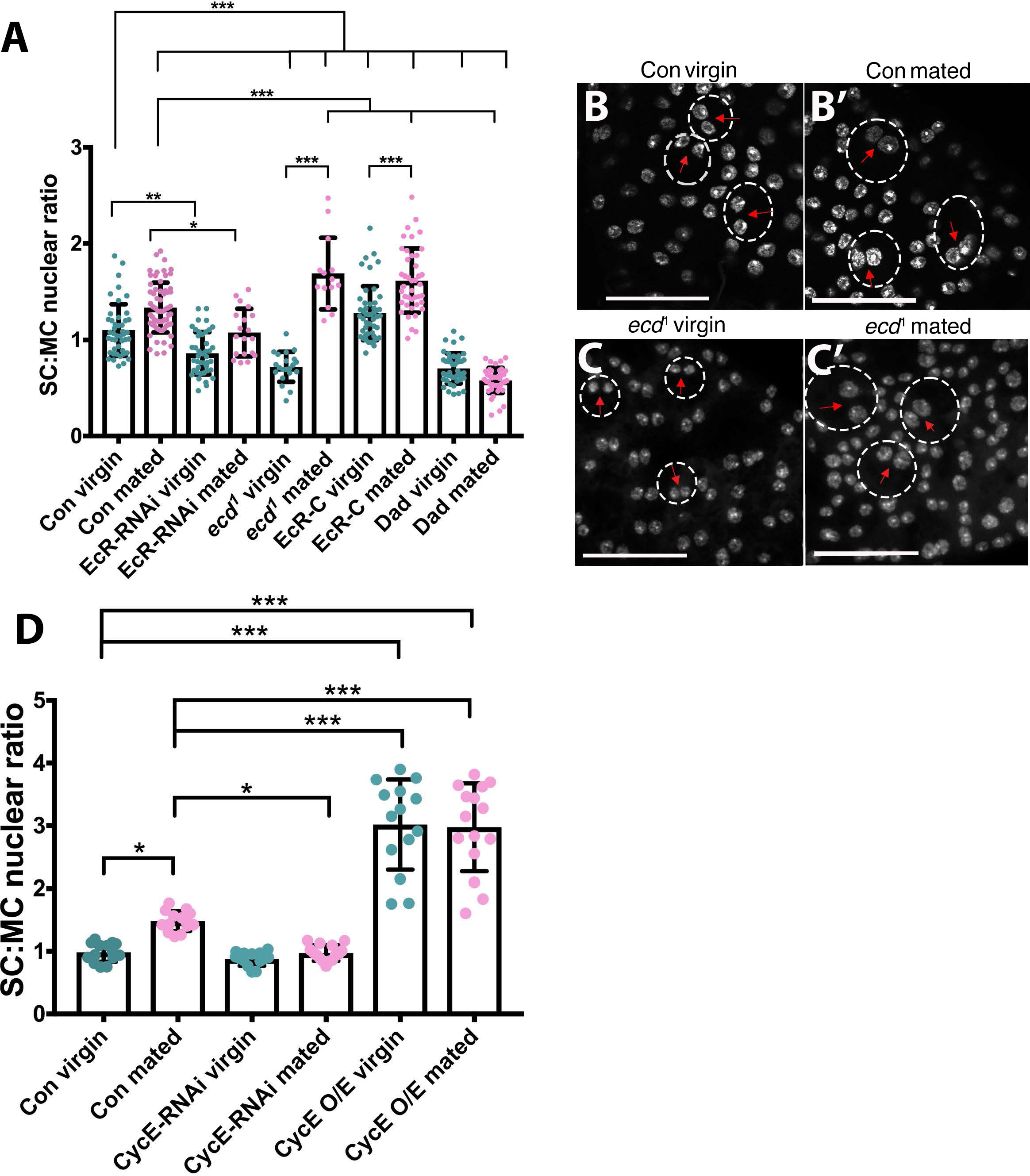
Mating induces hormone-independent, EcR-regulated SC nuclear growth. A. Histogram showing SC nuclear size relative to adjacent MC nuclei in 6-day-old males for control glands, glands expressing EcR-RNAi, EcR-C or the BMP antagonist Dad in SCs under esg^ts^F/O control, and *ecd*^*1*^ mutant glands. Mating induces nuclear growth in control glands (eg. see B), which is suppressed when EcR and BMP signalling is reduced, but enhanced by the *ecd*^*1*^ mutant (see C). B,B’, C,C’. SC nuclei (marked by red arrows; stained with DAPI) in *ecd*^*1*^ virgin males (C) are much smaller than controls (B), while SCs in mated *ecd*^*1*^ glands (C’) have increased nuclear growth. SCs from glands of 6-day-old males were identified by their characteristic morphology and their approximate outline is marked by a dashed circle. D. Histogram showing effects of RNAi-mediated knockdown of *cycE* and overexpression of *cycE* in SCs. Data in A and D analysed using one-way ANOVA and Tukey’s multiple comparisons test; *p<0.01, **p<0.001, ***p<0.0001. n=15. Scale bars, 50 μm.

### EcR protein levels are controlled post-transcriptionally by BMP signalling to regulate growth

Since EcR-B1 is the major isoform expressed by SCs, we examined the effect of overexpressing it in these cells under esg^ts^F/O control. Unexpectedly, SC nuclear size was not affected either by this treatment (Fig. 2C, F), or by overexpression of EcR-A or EcR-B2 (Fig. S1D, F, H). However, when we analysed EcR protein levels in these backgrounds, they appeared unchanged compared to controls (Figs. 1G, S2A, D and F), even though these constructs increase EcR levels when expressed in adjacent MCs (Fig. S3A, C, E and G). Since many other UAS-coupled transgenes can be overexpressed in SCs under esg^ts^F/O control, our data suggest that EcR levels are tightly controlled post-transcriptionally in these cells, so that increased *EcR* transcription has no obvious effect on receptor levels or growth.

Previous work has shown that SC growth is positively regulated by BMP signalling [9]. Since EcR signalling also promotes growth, we investigated the effect of BMP signalling on EcR protein expression. SC-specific expression of a constitutively active form of the Type I BMP receptor Thick veins (Tkv^Q199D^ or Tkv^ACT^) [45] induced increased levels of EcR protein, which was primarily localised in the nucleus (Fig. 4B). When BMP signalling was reduced in SC mosaics by inducing SC-specific knockdown of *Med*, encoding a downstream co-Smad transcription factor in the BMP signalling pathway, virtually no EcR protein was observed in knockdown cells (Fig. 4C). A similar BMP-dependent effect on EcR levels was not observed in main cells and main cell growth was also unaffected by Tkv^ACT^, EcR or combined Tkv^ACT^/EcR expression (Fig. S3). Taken together, these data indicate that BMP signalling is a key regulator of EcR protein levels specifically in SCs.

**Figure 4.**
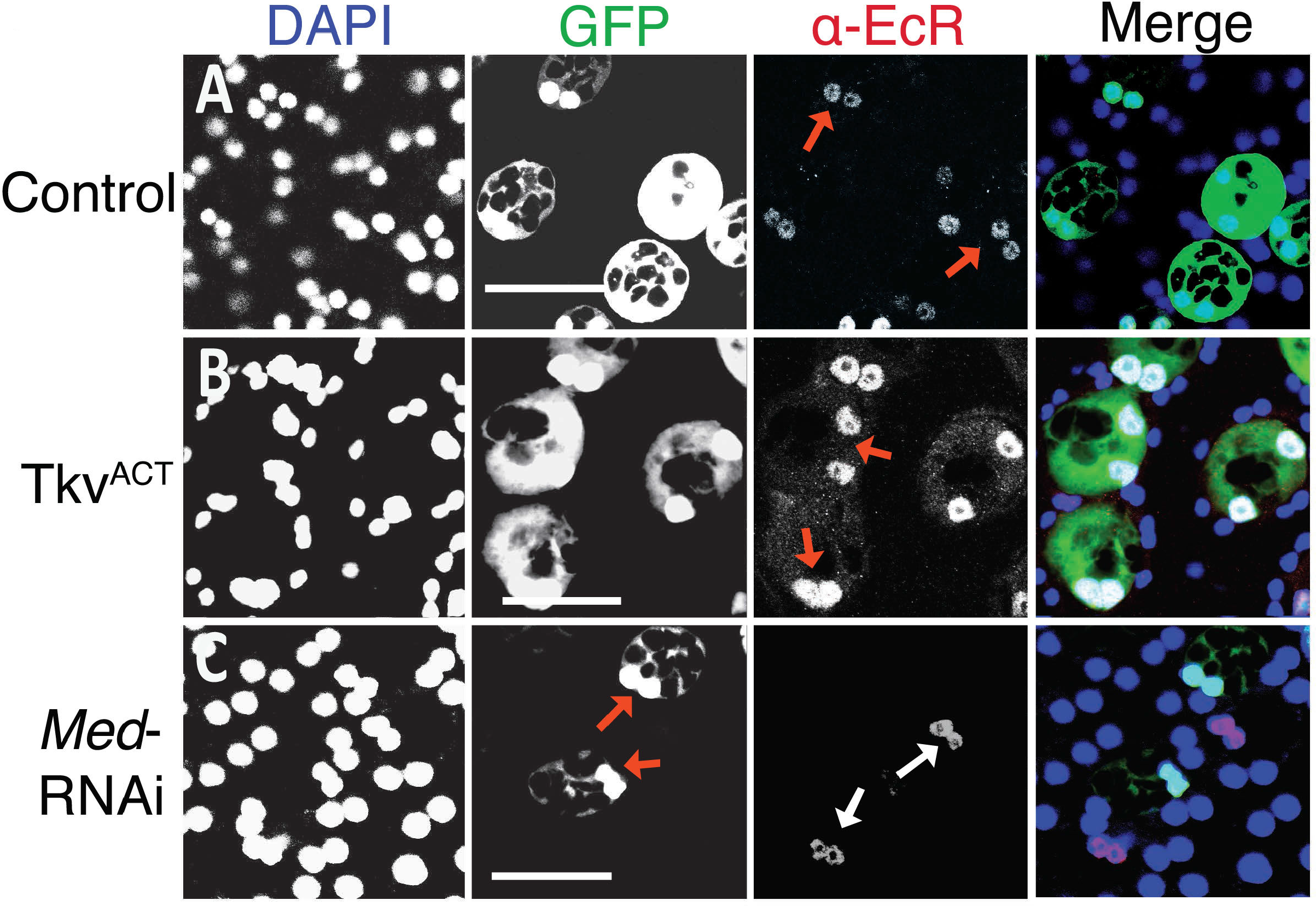
BMP signalling regulates levels of the EcR protein in SCs. Images show the AG epithelium dissected from 6-day-old virgin males expressing GFP and other transgenes under esg^ts^F/O control, and stained with a pan-EcR antibody. A, B. Upregulation of BMP signalling by SC-specific Tkv^ACT^ overexpression in adults results in increased expression of EcR (B) compared to control (A), with an enhanced nuclear signal (red arrows), and some cytosolic expression. C. RNAi knockdown of Medea in only some SCs, by activating the esg^ts^F/O driver system in 3-day-old adults, reduces BMP signalling in these cells and leads to a marked reduction in SC-specific EcR expression (nuclei marked with red arrows) after a further 6 days. EcR expression in SCs that do not express the RNAi construct is normal (white arrows). Scale bars, 50 μm.

To analyse the interaction between BMP and EcR signalling further, we next tested whether overexpressing EcR when BMP signalling is hyper-activated might further increase EcR protein levels and promote growth. Co-overexpression of Tkv^ACT^ and EcR-B1 in SCs resulted in a strong synergistic enhancement of growth (Figs. 5A-D, H). Similar synergistic growth effects were observed with both EcR-A and EcR-B2, even though overexpressing either EcR isoform in the absence of Tkv^ACT^ had no effect on nuclear size (Fig. S1A, C-H).

**Figure 5.**
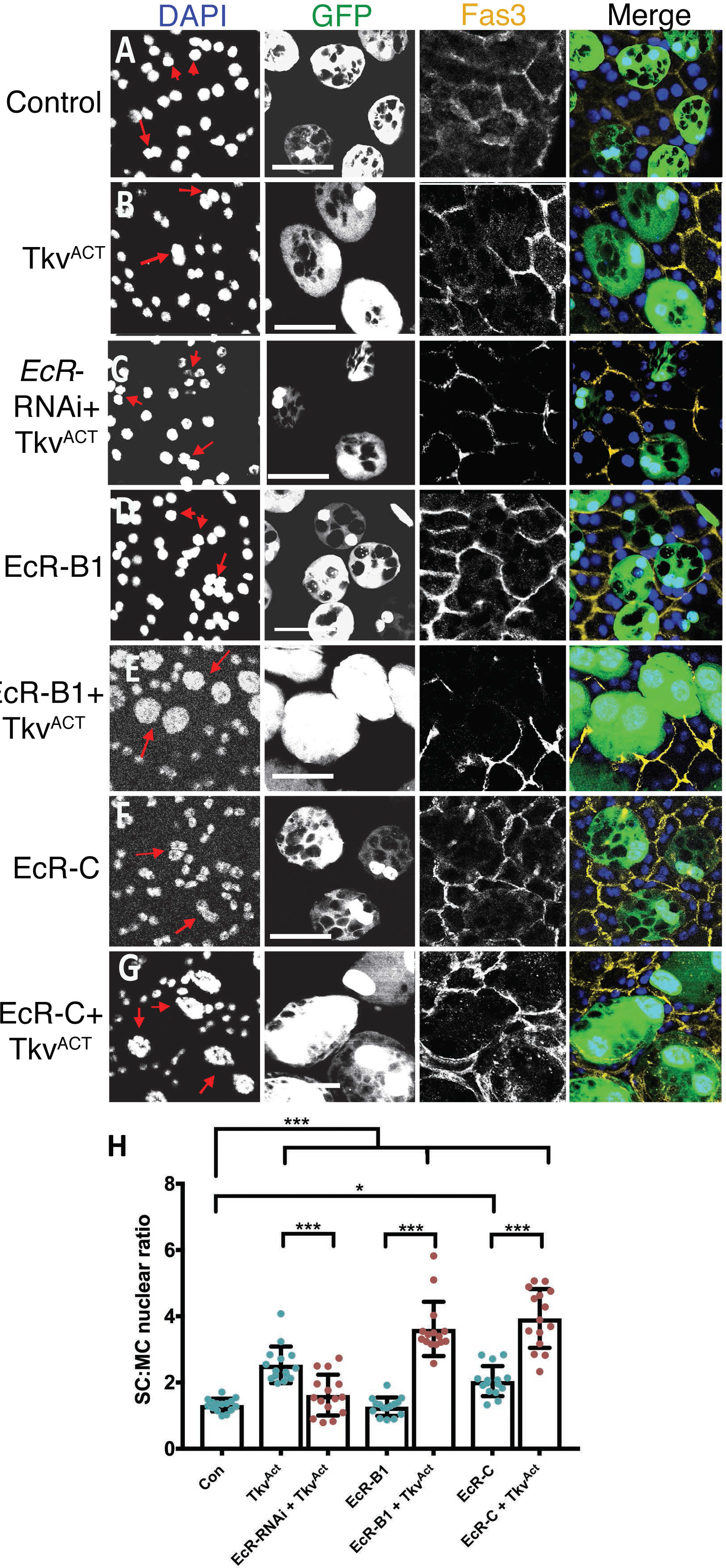
BMP signalling and the EcR synergise to regulate SC growth. Dissected AGs from 6-day-old virgin males expressing GFP and other transgenes under esg^ts^F/O control were stained with an antibody against Fas3 to mark the apical outlines of SCs and neighbouring MCs (yellow) and DAPI (blue nuclei). A-C. The nuclear growth induced by SC-specific expression of Tkv^ACT^ (B) is completely suppressed by co-expression of EcR-RNAi (C) to levels comparable with controls (A). D-G. Co-expression of Tkv^ACT^ (to upregulate BMP signalling) and EcR-B1 (E) or EcR-C (G) produces a synergistic enhancement of nuclear (and cell) growth in 6-day-old adults relative to the effects of Tkv^ACT^ (B), EcR-B1 (D) or EcR-C (F) alone. Note that some main cells are compressed between the giant co-expressing SCs. H. Histogram showing size of SC nuclei relative to MC nuclei in AGs where SCs are expressing Tkv and EcR transgenes. Selected SC nuclei are marked with red arrows. Data analysed using one-way ANOVA and Tukey’s multiple comparisons test *p<0.02, ***p<0.0001. n=15 Scale bars, 50 μm.

In addition to these dramatic growth effects, very high levels of EcR expression were observed in SCs co-expressing Tkv^ACT^ and any of the EcR isoforms (Fig. 6A-D and S2A, D-G). The highest EcR levels were observed in nuclei, but all co-expressing cells also had detectable cytosolic EcR, which was highly elevated in some cells (Fig. 6D and F, S2E and G). We conclude that BMP signalling primarily controls the levels of EcR protein expression post-transcriptionally in SCs.

**Figure 6.**
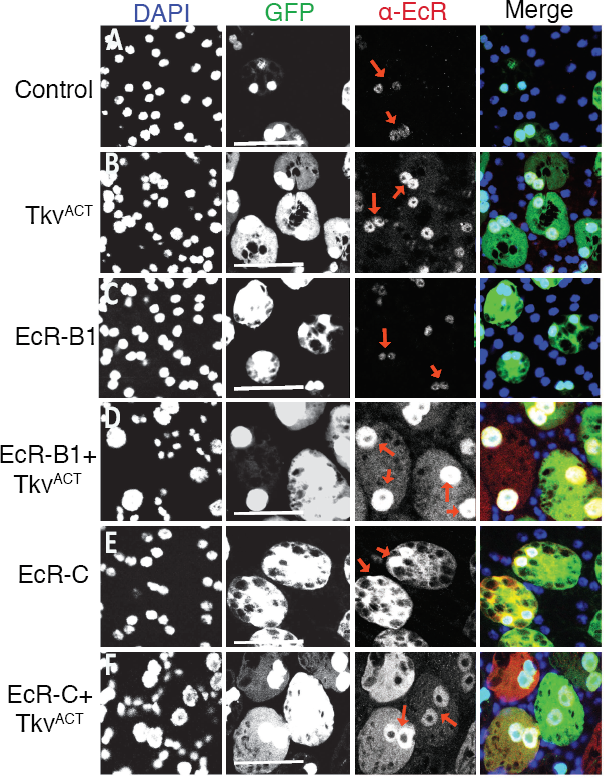
BMP signalling regulates EcR levels post-transcriptionally in SCs, probably via the EcR N-terminal domain. Dissected AGs from 6-day-old virgin males expressing GFP and other transgenes under esg^ts^F/O control were stained with a pan-EcR antibody (red arrows) and DAPI (blue nuclei). A, B. Nuclear EcR levels in SCs are elevated when BMP signalling is increased upon expression of Tkv^ACT^ in SCs (B) compared to controls (A). C, D. Nuclear EcR levels are unaffected by overexpression of EcR-B1 alone (C), but levels of nuclear protein are highly upregulated when co-expressed with Tkv^ACT^ (D). Some cytosolic EcR is also present. E, F. By contrast, overexpression of the N-terminally truncated EcR protein, EcR-C, leads to accumulation of nuclear and cytosolic EcR (E). Co-expression with Tkv^ACT^ appears to increase the ratio of nuclear to cytosolic EcR in some cells (F). Scale bars = 50 μm.

In light of this strong dependence of growth-regulatory EcR protein levels on BMP signalling, we tested whether BMP-dependent growth in SCs is mediated through EcR signalling by co-expressing Tkv^ACT^ with pan-*EcR*-RNAi. Tkv^ACT^-induced growth was strongly suppressed, and the resulting SC nuclei were not significantly different in size from wild type controls (Fig. 5A, B, G and H), indicating that BMP-dependent SC growth requires the presence of the EcR.

### The unique AF1-containing N-terminal domain of each EcR protein isoform appears to be involved in its SC-specific post-transcriptional regulation by BMPs

Previous *in vitro* studies have revealed that when the *Drosophila* EcR-A and EcR-B1 isoforms are expressed in Chinese Hamster Ovary (CHO) cells, the different N-terminal domains (NTD) of these proteins, which include the AF1 domain, one of the two transcriptional activation domains in these receptors, partially destabilise the proteins through a ubiquitination-dependent mechanism [46]. Although these experiments were performed in a heterologous system and did not reveal similar regulation for the EcR-B2 isoform, which has a much shorter AF1 domain, we tested whether the NTD plays a role in SC-specific, BMP-dependent control of EcR protein levels. We expressed EcR-C, an artificial isoform of the protein in which the NTD sequence has been deleted, in SCs. This protein only contains sequences common to all isoforms [47] and in other cell types it usually has reduced activity compared to native forms of EcR when overexpressed.

Unlike the endogenously expressed isoforms of EcR, overexpression of EcR-C promoted growth of SCs (Fig. 2D, F) and produced high levels of EcR protein in the nuclei and cytosol of SCs when expressed alone (Fig. 6E). The most likely explanation of our data, given the known role of EcR NTD sequences in protein stability [46], is that BMP signalling activity regulates levels of different EcR isoforms via a genetic interaction with each of their unique NTDs. The growth effects of EcR-C were further enhanced by co-expression with Tkv^ACT^ in virgin males (Fig. 5F and H), indicating that BMP signalling affects EcR activity via more than one mechanism. Indeed, relative levels of nuclear versus cytosolic EcR-C appeared to be increased by Tkv^ACT^ overexpression in many, but not all, SCs, suggesting that BMP signalling might also regulate the nuclear trafficking of EcR (Fig. 6F). When males, which are overexpressing EcR-C in SCs, were mated, some additional SC growth was observed compared to virgins (Fig. 3A), perhaps because of the associated increase in BMP signalling.

A post-transcriptional interaction between the EcR and BMP signalling pathways has not previously been reported in *Drosophila*. It was not observed in main cells (Fig. S3I, J). We conclude that BMP signalling controls EcR levels and EcR signalling in SCs post-transcriptionally via a cell type-specific interaction that appears to partly involve the EcR NTD.

### Mating drives synthesis of new DNA in SCs, which is regulated by BMP-dependent, but hormone-independent, EcR signalling

In *Drosophila*, increased nuclear and cell size is often associated with endoreplication, which increases gene expression through elevated gene copy number [48]. At eclosion, both SCs and MCs have two large nuclei, each estimated to be tetraploid [49]. Previously we were unable to detect endoreplication in adult SCs of mated flies fed with the nucleotide analogue bromodeoxyuridine (BrdU) [9]. However, we reasoned that this might be explained by poor penetration of the anti-BrdU antibody in AGs, and therefore repeated these experiments by feeding males with 5-ethynyl-2′-deoxyuridine (EdU), which can be detected chemically, throughout adulthood. While EdU uptake was rarely observed in SCs of 6-day-old virgin males, approximately 25% of SCs from multiply-mated males incorporated EdU, indicating that new DNA synthesis was occurring in a subset of these cells (Fig 7A, B and G). No new DNA synthesis was observed in the MCs of either virgin or mated glands (Fig 7A and B).

**Figure 7.**
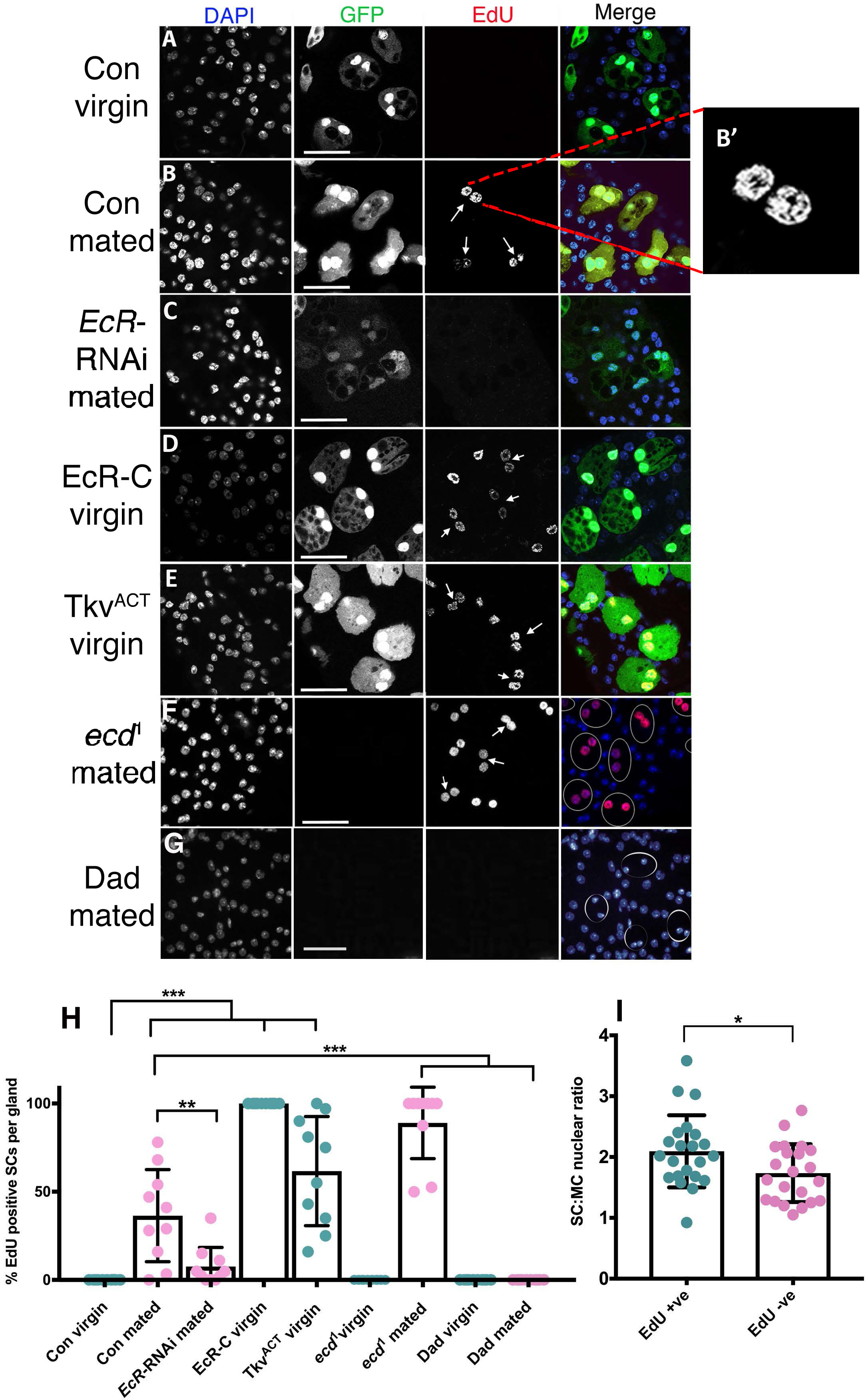
Hormone-independent, EcR-mediated endoreplication of SC DNA is stimulated by mating. Males expressing GFP and other transgenes under esg^ts^F/O control or *ecd*^*1*^ mutants were cultured on EdU-containing food post-eclosion, dissected at 6 days, and their AGs probed for EdU uptake to assess DNA replication and stained with DAPI. A, B. EdU was incorporated in about 30% of SCs after mating (B; white arrows depict SC with EdU uptake), but not in virgins (A). Inset in B shows high magnification view of single SC. C. SC-specific expression of EcR-RNAi blocks EdU incorporation in mated males. D, E. Expression of EcR-C (D) or Tkv^ACT^ (E) in SCs promotes EdU incorporation in SCs from virgin males. F. Almost all SCs in *ecd*^*1*^ males incorporate EdU in their nuclei after mating. G. Dad-expressing SCs do not incorporate EdU after mating (note weak GFP expression in these cells is masked following the EdU staining procedure). H. Histogram showing EdU incorporation into SC nuclei in different genetic backgrounds. I. Histogram showing SC:MC nuclear size ratio for EdU-positive and -negative SCs from esg^ts^F/O control mated males. One-way ANOVA, Dunnett’s multiple comparisons test. *p<0.0001, n=15. Scale bars, 70μm.

Interestingly, SC nuclei that take up EdU in mated glands were larger than the EdU-negative nuclei (Fig 7H), demonstrating that part of the increase in SC nuclear size is a consequence of new DNA synthesis. Furthermore, EdU incorporation was distributed across all parts of the nucleus (Fig 7B’), suggesting that it is not the result of focal gene amplification, as is seen for chorion genes in ovarian follicle cell nuclei [50]. To further assess whether genome endoreplication is responsible for mating-dependent growth, we tested the effect of overexpressing and knocking down the G1/S cyclin, Cyclin E (CycE) in SCs, which is required for endoreplication in other *Drosophila* cell types [48]. While knockdown of *cycE* had no significant effect on nuclear growth in virgin males, it inhibited the additional growth in mated males (Fig. 3D). Furthermore, overexpression of *CycE* in virgin males stimulated nuclear growth, but this was not enhanced by mating (Fig 3D). Consistent with these findings, studies of endoreplication in the salivary gland have suggested that constant overexpression of CycE can drive one cycle of endoreplication, but does not permit further rounds [51].

Given that both BMP and EcR signalling modulate SC nuclear growth in mated males, we tested whether these pathways regulate DNA synthesis in adult SCs. In complete contrast to controls, the majority of SCs expressing Tkv^ACT^ in adult virgin males typically incorporated EdU over 6 days (Fig 7F and G). Furthermore, all SCs expressing the EcR-C construct contained nuclear EdU (Fig 7E and G). EdU uptake was significantly suppressed in glands from multiply-mated males expressing EcR-RNAi or the BMP antagonist Dad in SCs (Fig 7D, G and H).

Finally, the number of SCs incorporating EdU in *ecd*^*1*^ males shifted to the non-permissive temperature after eclosion was assessed. This genetic manipulation had no effect on EdU incorporation in virgin males. By contrast, incorporation in multiply-mated males was significantly increased compared to controls, with positive staining in virtually all SCs, demonstrating that mating-induced, EcR-mediated endoreplication is not hormone-dependent, and indeed, may be partially inhibited by ecdysone (Fig. 7C and G). Taken together, these data indicate that both BMP and EcR signalling act in SCs of mated males to promote synthesis of new DNA. This endoreplication explains much of the additional nuclear growth in SCs after mating (Fig. 3A), but unlike growth in virgin males, this process is ecdysone-independent.

## Discussion

Secretory cells in both the mammalian prostate and the fly accessory gland have unusual growth properties in adults. Androgens play a central role in regulating growth and proliferation in the prostate, potentially linking nutrition and sexual activity to adult glandular activity [22,23], as well as driving maturation during puberty. In advanced prostate cancer, when tumour cells can become resistant to anti-androgen treatment, they frequently grow via a hormone-independent, androgen receptor-driven process that remains incompletely understood [24].

Here we demonstrate that SC nuclear growth in virgin male flies also involves hormone-dependent steroid receptor activity. This activity is modulated by local autocrine BMP signals via a novel post-transcriptional mechanism. After mating, EcR-dependent nuclear growth becomes primarily hormone-independent and requires CycE-driven endoreplication. As we discuss below, these regulatory mechanisms potentially allow SC growth and secretion to adapt to socio-sexual experience and mating.

### BMP signalling tightly regulates EcR levels to control SC growth

Our findings that BMP signalling is required in SCs for them to express detectable levels of a specific EcR isoform, EcR-B1, and that knocking down EcR expression blocks BMP-stimulated growth, strongly indicate that EcR signalling is a primary mediator of BMP-dependent growth regulatory effects in these cells.

The BMP/EcR post-transcriptional interaction appears highly cell type-specific in flies. It cannot be induced in main cells of the AG and has not been reported in multiple tissue types during *Drosophila* development. Signalling by Activins, members of another class of TGF-β ligands, is required for EcR-B1 expression during neuronal remodelling of the fly brain at metamorphosis [52]. However, the effects of losing the Type I Activin receptor Baboon can be overcome by GAL4/UAS-driven expression of EcR-B1, suggesting that SC-like post-transcriptional control of EcR-B1 is not involved [52]. Regulation of EcR-B1 expression by TGF-β/BMP signalling has also been reported in larval motoneurons as they dismantle during metamorphosis, but again post-transcriptional control has not been implicated [53].

Each of the three normal isoforms of EcR has a unique N-terminal domain, which includes a so-called AF1 transcriptional activation domain. The sequences encoding these NTDs appear to be essential for normal BMP-dependent regulation of EcR levels in SCs, because their absence in the EcR-C protein leads to partial evasion of this control. The transcript sequences encoding the normal EcR isoforms and EcR-C are all identical except in the 5’ regions that encode the isoform-specific N-terminal domains. It is possible that these different 5’ sequences are all independently targeted by a mechanism that mediates BMP-dependent control of EcR protein levels, but cannot affect EcR-C. However, it seems much more likely that the regulation is post-translational, particularly since the NTDs of both EcR-A and EcR-B1 are involved in degradative mechanisms that control EcR protein levels [46].

### Hormone-independent non-canonical EcR signalling in SCs is activated by mating and specifically regulates endoreplication

Not only is the EcR regulated via a unique post-transcriptional mechanism in SCs, it also has an unusual and complex mode of action and target specificity. Although Usp, the well-characterised binding partner of EcR during development, is expressed selectively in the nuclei of SCs, *Usp* knockdown does not alter SC nuclear size or EcR localisation. Usp-independent EcR signalling has been reported previously in one developmental scenario in larvae [54]; the mechanisms involved have not been characterised, although the authors propose that the EcR may act as a homodimer or bind with an alternative partner. Furthermore, recent work has shown that EcR expression is functionally important in development of the adult AG epithelium, but does not require Usp [34]. These authors did not identify the cells involved or the precise cellular defects.

We have screened for expression of several of the known target genes of EcR in development, such as *Broad* and *Eip74EF*, using well-characterised antibodies [55] and specific gene traps, but have not been able to identify any downstream targets of the EcR in SCs. Cell type-specific analysis of genomic EcR binding sites or the SC transcriptome will be required to unravel the genetic programme controlled by this receptor in these cells.

The other unique feature of EcR signalling in SCs is that mating alters its downstream effects. EcR-regulated nuclear and cell growth in virgins occurs independently of DNA replication, the former potentially reflecting decondensation of chromatin. But after mating, a subset of cells activates EcR-dependent endoreplication. This is responsible for much of the additional SC growth observed after mating, because the nuclei of these cells are larger and *cycE* knockdown specifically suppresses mating-dependent growth. The changes can be phenocopied in virgin males by activation of BMP signalling or overexpression of EcR-C in SCs. Most importantly, this EcR-regulated effect is hormone-independent, unlike growth in SCs of virgins. Remarkably, in most AGs from mated *ecd*^*1*^ males, the majority of SCs endoreplicate their genomes, suggesting ecdysone normally suppresses this process. In this context, the EcR could either be acting as a repressor of gene transcription, whose repression is released by ecdysone [39], or it may only bind to the targets, which it activates, in the absence of ligand. Analysis of mating-specific, EcR genomic binding sites will be required to distinguish these two hypotheses.

Hyper-activated BMP signalling may be an important trigger for endoreplication after mating. We have previously shown that only about 30% of SCs increase BMP signalling detectably following copulation [13], mirroring the proportion of endoreplicating cells under these conditions. However, we have yet to develop a robust protocol with which we can co-detect EdU and the BMP transcriptional target, P-Mad, to confirm that these two populations are the same. Furthermore, since genetically activating BMP signalling in SCs does not induce endoreplication in all cells, there is probably a second, as yet unidentified, mechanism that modulates the number of endoreplicating cells.

### The EcR provides a link between adult socio-sexual behaviour and accessory gland function

Other studies have shown that whole animal 20-hydroxyecdysone (20-HE) titres increase in male flies exposed to previously mated females [27] and that EcR signalling activity in the fly brain is required for normal courtship behaviours [27,29]. Furthermore, application of topical 20E to males exposed to females, which have an experimentally sealed ovipositor that prevents mating, rescues the reduction in AG secretory activity exhibited by these animals [33], strongly suggesting that the overall activity of the gland can be influenced by this hormone.

We propose that direct effects of ecdysone on SCs, cells which have an important role in AG reproductive function [9–11] and can affect the normal processing of main cell products like Ovulin [11], provide one route by which this hormone can alter the activity of the entire gland. Indeed, Sitnik et al [56] have presented evidence that blocking the normal development of SCs in a specific *Abd-B* mutant may have indirect effects on the transcriptional programme of MCs, further supporting the idea that SCs can co-ordinate functions of both epithelial cell types in the AG.

After mating, EcR-mediated growth of some SCs becomes hormone-independent and involves endoreplication. Endoreplication is employed to promote high levels of transcriptional activity and secretion in a range of organisms from mammals to plants [48], for example in the salivary glands and follicle cells of the egg chamber in *Drosophila*. New DNA synthesis in SCs is therefore likely to boost transcription in this highly specialised, secretory cell type to replenish SC products released from the AG during mating, in preparation for subsequent matings.

Such regulation has important physiological implications. Ecdysone levels and SC growth are reduced in virgin males, particularly in the absence of females [27], thus conserving resources. The hormone-independent endoreplication mechanism allows such flies to rapidly upregulate SC activity following mating, using the EcR (Figure 8). Indeed, since *ecd*^*1*^ mutant males exhibit an enhanced level of endoreplication after mating, virgin males with the lowest ecdysone levels and therefore the least SC growth, could respond particularly strongly to these post-mating, hormone-independent signals.

**Figure 8.**
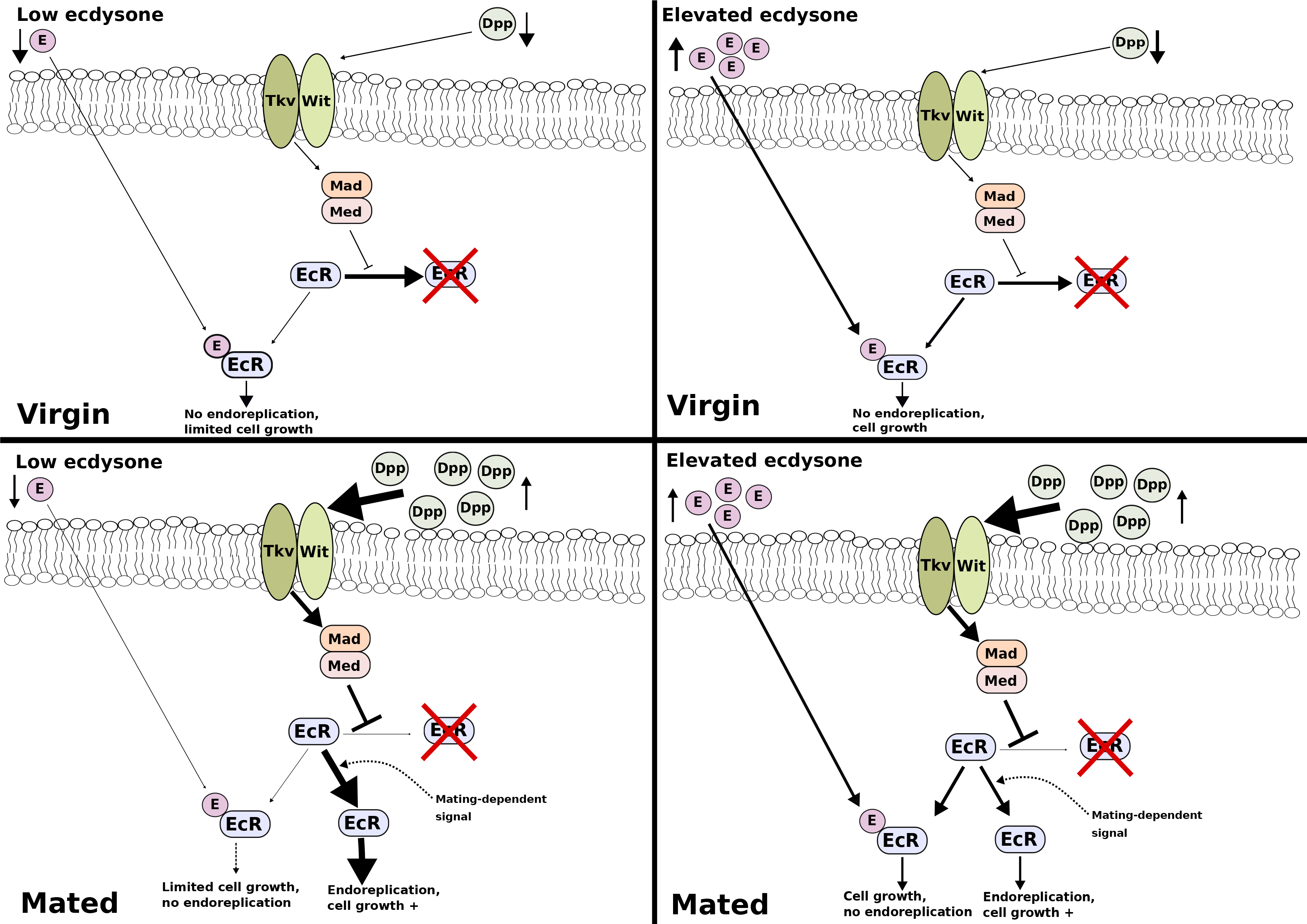
Proposed model explaining physiological basis of hormone-dependent and -independent, EcR-mediated SC growth. Our data reveal that EcR-dependent growth of SCs is differentially modulated by the presence of ecdysone (E), according to mating status. An E-EcR complex, which does not require Usp, appears to be necessary for normal SC nuclear growth observed in virgin males. This form of growth does not involve endoreplication. Growth is restricted in virgin males with low E titres, whilst virgin males with higher E titres, such as those in contact with pre-mated females, should have larger SCs and presumably more biosynthetic and secretory activity. In mated males, SC growth is enhanced, at least in part due to new DNA synthesis. Growth and the proportion of SCs that endoreplicate their genome is greatly enhanced in mated *ecd*^*1*^ males, but suppressed in *EcR*-RNAi-expressing SCs, indicating that mating-induced growth and endoreplication occurs via a hormone-independent, EcR-mediated mechanism. This is stimulated by elevated BMP signalling induced by autocrine BMP ligand Dpp through the heterodimeric Tkv/Wit receptor, which appears to stabilise EcR [13]. In mated males, EcR that is not bound to E must either repress or activate a subset of genes that are not EcR targets in virgin males, hence inducing endoreplication, and this is suppressed by E. Direct or indirect targets include cell cycle regulators like *cycE*. In mated males with no ecdysone, more SCs switch to endoreplication-dependent growth. This mechanism permits small SCs from males with low E levels (for example, because they have been isolated from females) to grow more rapidly in response to mating, when compared to SCs from males with higher E titres, which are already enlarged in virgins.

### Growth in both the prostate cells and the fly secondary cells is regulated by BMPs and different forms of steroid receptor signalling

We initially tested the function of EcR signalling in SCs because of the critical role of androgens and AR signalling in normal and tumorigenic prostate epithelial growth. We have uncovered a clear parallel between humans and flies: the receptor for a steroid hormone, which is regulated by socio-sexual experience and environment, is involved in controlling growth of secretory cells in both male glands, and can function by hormone-dependent and ‒independent mechanisms. In flies, this switch plays a physiological role, while in prostate, it has only been observed to date in cancer.

Like steroids, BMP signalling has also been implicated in prostate growth and metastasis [15–17]. However, its effects are complex, because different BMP ligands, which signal through alternative pathways, can have opposite effects. In prostate cancer, BMP signalling has been implicated in the androgen-independent AR signalling associated with castration resistance [57–59], though its mode of action remains unclear. Our findings suggest links between BMPs and steroid receptors in the male reproductive system of the fly that switch the EcR to a hormone-independent mode under physiological conditions. The parallels in the mechanisms involved now require further investigation in both flies and humans with particular focus on defining the cellular conditions under which BMP-induced, hormone-independent signalling is activated.

## Materials and Methods

### Fly strains and culture

The following fly strains (obtained from the Bloomington Stock Centre, except where noted) were employed: esg^ts^F/O (*w; esg-GAL4, UAS-GFPnls; act>CD2>GAL4, UAS-FLP*; gift from B. Edgar) [35], *UAS-EcR-RNAi* (TRiP.JF02538) [60], *UAS-EcR-B1*, *UAS-EcR-A*, *UAS-EcR-B2, UAS-EcR-C* [47,61], *UAS-EcR-B1-RNAi*, *UAS-EcR-A-RNAi* [38], *UAS-Tkv*^*Q199D*^ [45], *UAS-Med-RNAi* [62], *UAS-Usp-RNAi* (TRiP.JF02546), *UAS-ry-RNAi* (TRiP.44106), and *Acp26Aa-GAL*4 (gift from S. Goodwin; [63]. TRiP UAS-RNAi lines are described in [64]. Flies were fed on standard cornmeal agar medium. No dried yeast was added to the vials.

### Fly Genetics

To express UAS-transgenes in adult SCs under esg^ts^F/O control [9], fly crosses were initially cultured at a non-permissive temperature, 18°C or 25°C (*w*^*1118*^ and *UAS-ry-RNAi* flies were used in control crosses with the esg^ts^F/O strain). Newly eclosed, virgin males of the appropriate genotype were selected, separated from females and transferred to 28.5°C immediately. All SCs that induce FLP-mediated recombination of the *act>CD2>GAL4* construct continue to express GFP. Mosaic experiments were performed by delaying the temperature shift until day 3 of adult life, with dissection of adults at approximately 9 days post-eclosion [9]. Expression of *esg-GAL4* is gradually lost in some adult SCs, so delaying the temperature shift results in *act>CD2>GAL4* recombination in a subset of SCs. For nuclear size measurements and growth analysis, males were typically dissected at 6 days.

### Immunohistochemistry and imaging

This followed previously published methods [9,14]. Flies were anaesthetised using CO2 and dissected with fine forceps in 4% paraformaldehyde dissolved in PBS. Dissected AGs were transferred to Eppendorf tubes, fixed for 20 min at 22°C and then washed 6 × 10 min in 1 ml PBST [1 × PBS, 0.3% Triton X-100 (Sigma-Aldrich)]. Anti-Fas3 [65], anti-pan-EcR, anti-EcR-A, and anti-EcR-B1 [36] antibodies were all obtained as supernatants from the Developmental Studies Hybridoma Bank, Iowa and diluted 1 in 10 in PBSTG (PBST, 10% goat serum). Mouse anti-Usp antibody (1:100 dilution) was a kind gift from the Kafatos lab [44] and the rabbit anti-ANCE antibody (1 in 200) was kindly provided by E. Isaac [66]. Glands were incubated overnight at 4°C in primary antibody. They were then washed for 6 × 10 min in PBST before incubation with either Cy3- or Cy5-conjugated donkey anti-mouse secondary antibody (Jackson laboratories) used at a dilution of 1 in 400 for 2 hours at room temperature. Glands were further washed in PBST for 6 × 10 min, before mounting on slides using DAPI-containing Vectashield (Vector Laboratories). Imaging of glands was performed using a Zeiss Axioplan 2 scanning confocal microscope with a LSM510 laser module or a Zeiss 880 Airyscan system. Nuclear areas were measured using Axiovision freeware (Zeiss) as previously described (Leiblich et al., 2012).

### Detection of DNA replication using EdU

To detect DNA replication using the thymidine analogue, 5-ethynyl-2’-deoxyuridine (EdU), a Click-iT^®^ EdU imaging kit (Invitrogen) was used [67]. Adult flies were maintained on medium containing 0.2 mM EdU (ThermoFisher) from eclosion until dissection at 6 days. The EdU-containing medium was prepared by mixing standard yeast-cornmeal agar medium with a 10 mM stock solution (diluted in PBS, per manufacturer’s instruction). To detect EdU incorporation, dissected AGs were fixed in paraformaldehyde and processed as for immunohistochemistry. The Click-iT^®^ reaction mix was prepared following the manufacturer’s instructions, using a 1:400 dilution of 2mM stock of azide-fluor 555 (Sigma Aldrich) dissolved in DMSO. To label DNA in the sample, 200 μl of the reaction mix was added to the vials and left to incubate for 30 minutes at 20°C, away from light. Glands were washed three times in 200 μl PBST and then resuspended in 200 μl PBS, before mounting on coverslips using DAPI-containing mounting medium.

### Statistical analyses

We compared the mean SC:MC nuclear area across genotypes and controls. Having confirmed the data were normally distributed by the Shapiro-Wilk test, we used one-way analysis of variance (ANOVA) and Tukey’s multiple-comparison post-test to identify significant changes. Differences were deemed significant at a P value of <0.05. Statistical analyses were performed using GraphPad Prism 7.0, GraphPad Software, La Jolle California USA, www.graphpad.com. Identical statistical analyses were performed to compare the mean proportion of SCs incorporating EdU across genotypes with control glands.

## Acknowledgements

We thank Elwyn Isaac, Fotis Kefatos, Bruce Edgar, and Stephen Goodwin for stocks and reagents; we are grateful to the Bloomington *Drosophila* Stock Center for flies and to the Developmental Studies Hybridoma Bank (DSHB), Iowa for antibodies. Some microscopy was undertaken in the Micron Oxford Advanced Bioimaging Unit.

## Author contributions

AL, JEH, AS, DCIG, FCH, and CW conceptualised and planned experiments; AL, JEEUH, AS, CG, SR, SMW, and PM performed experiments; AL, JEH, AS, and CW analysed and evaluated the data; AL and CW wrote the manuscript; all authors read the manuscript.

## Supplementary Figures

**Figure S1.**
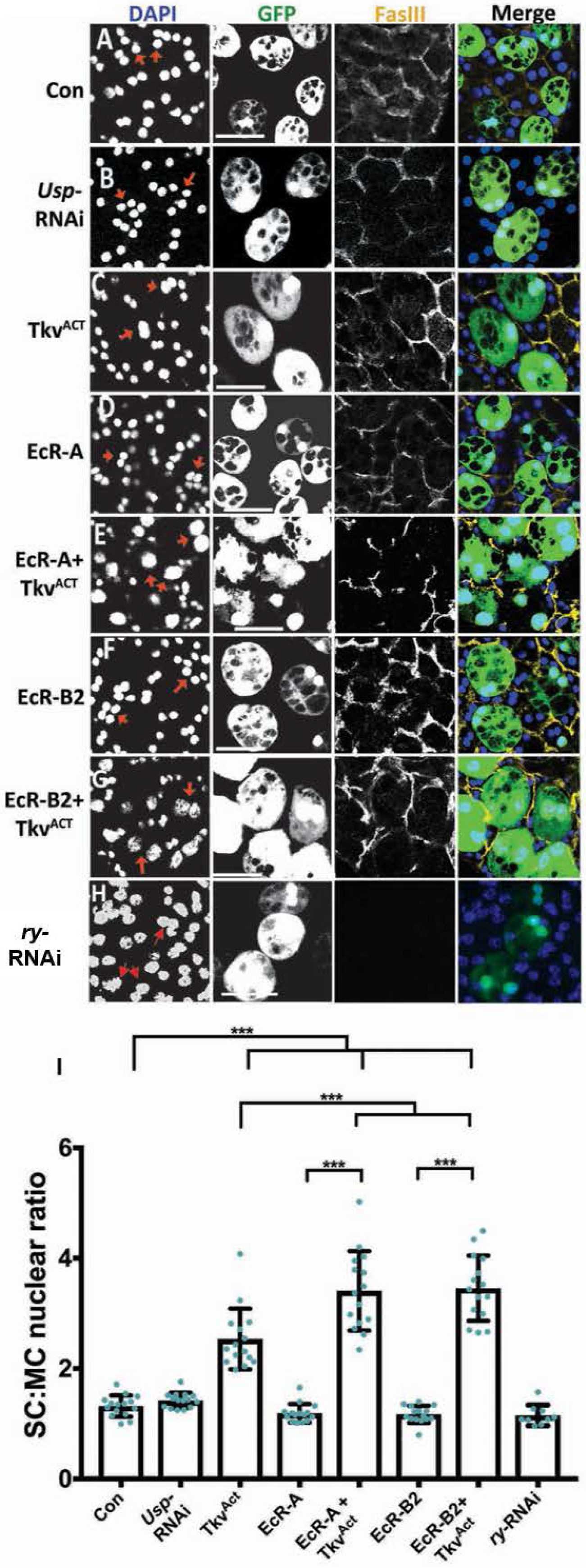
BMP signalling and the EcR synergise to regulate SC growth. Dissected accessory glands from 6-day-old males were stained with an antibody against Fasciclin3 to mark the apical outlines of SCs and neighbouring MCs (yellow) and with DAPI (blue nuclei). Selected SC nuclei are marked with red arrows and express GFP and other transgenes under esg^ts^F/O control. A, B. RNAi-mediated knockdown of *Usp* has no effect on SC nuclear growth (B) compared to control (A). D-G. Over-expression of the ‒A (D) and ‒B2 (F) isoforms of EcR has no effect on SC nuclear growth, but co-expression of these isoforms with Tkv^ACT^ synergistically promotes growth (E, G). H. RNAi-mediated knockdown of a control gene, *ry*, had no effect on growth. I. Histogram showing size of SC nuclei relative to MC nuclei in AGs where SCs express different transgenes as above. Significance was assessed by two-way ANOVA. ***p<0.0001, n=15. Scale bars, 60 μm.

**Figure S2.**
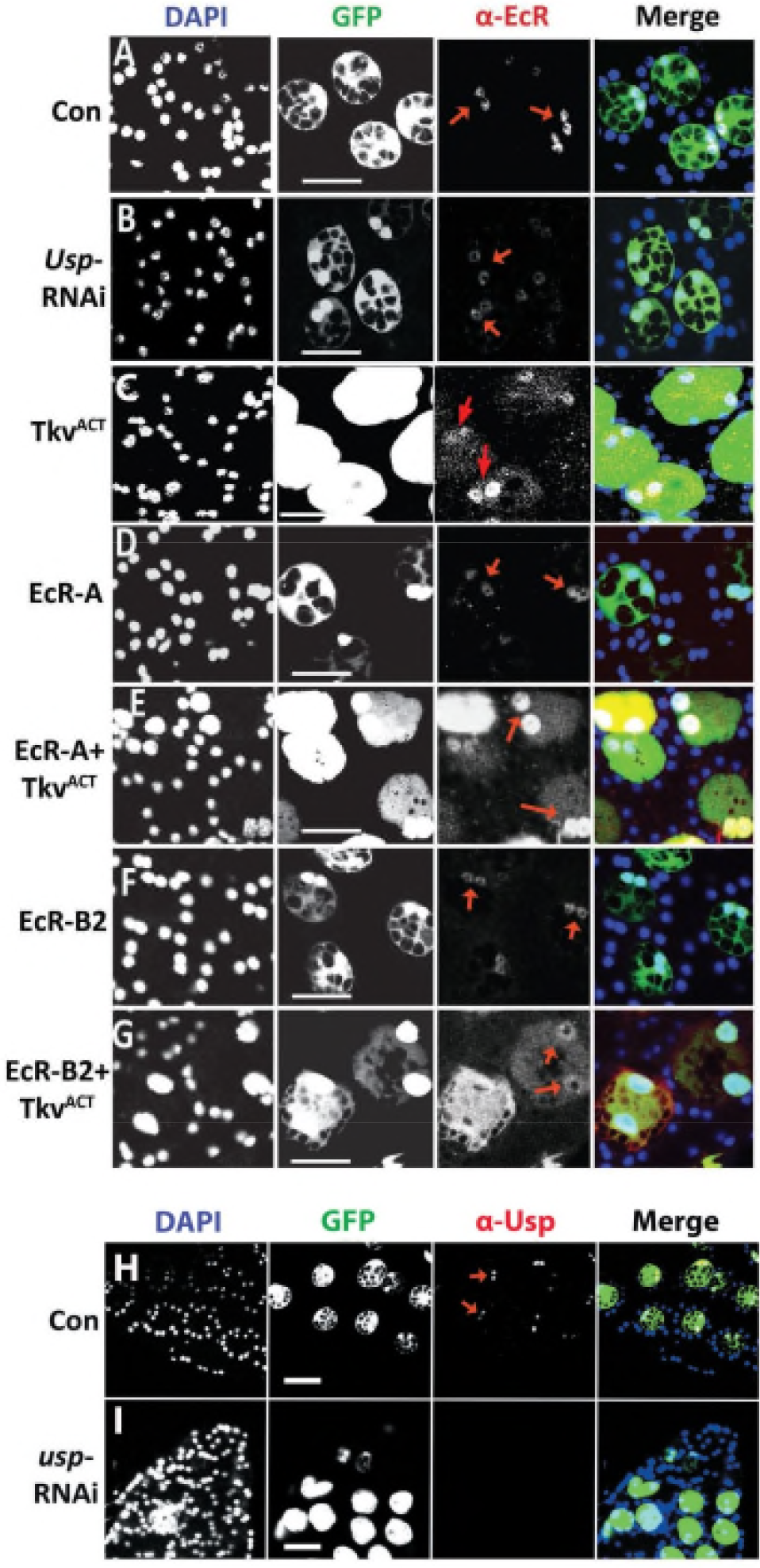
BMP signalling, but not Usp, regulates levels of the EcR protein in SCs. Images show the AG epithelium dissected from 6-day-old virgin males expressing nuclear GFP and other transgenes under esg^ts^F/O control, and stained with a pan-EcR antibody (A-G) or anti-Usp antibody (H,I). Nuclei are stained with DAPI (blue). Selected SC nuclei are marked with red arrows. A, B. *Usp* knockdown (B) has no effect on EcR expression compared to control (A). C-G. Overexpression of EcR-A (D) or EcR-B2 (F) does not appear to significantly alter EcR expression compared to controls (A). Co-expression of these isoforms with Tkv^ACT^ in SCs (E and G respectively) increases EcR expression in SCs compared to controls (A) and SCs expressing Tkv^ACT^ alone (C). H,I Immunostaining with an antibody that recognises Usp reveals expression in the nuclei of control SCs (H), but absence of expression in the nuclei of SCs expressing an RNAi targeting *Usp* (I). Scale bars, 60 μm (A-G), 120 μm (H, I).

**Figure S3.**
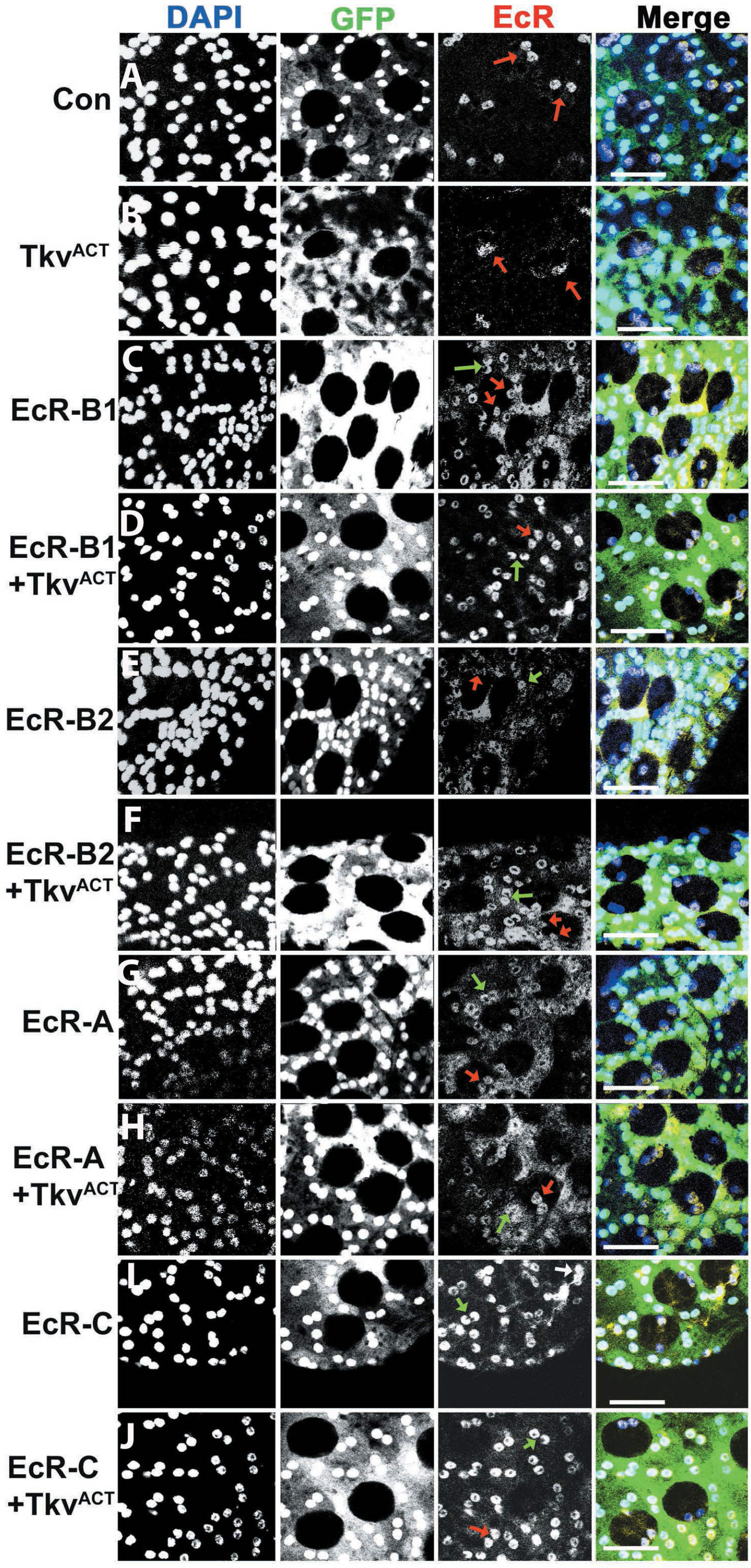
BMP signalling does not regulate levels of the EcR protein in main cells. Images show the AG epithelium dissected from 6-day-old virgin males expressing nuclear GFP and other transgenes in main cells under Acp26Aa-GAL4 control, and stained with a pan-EcR antibody. Nuclei are stained with DAPI (blue). A, B. Expression of Tkv^ACT^ in main cells (B), which do not normally express EcR (see control cells in A) does not affect EcR levels. C-J. Expression of EcR-B1 (C), -B2 (E), -A (G), and ‒C (I) in main cells leads to accumulation of EcR in these cells, in contrast to SCs. Co-expression with Tkv^ACT^ does not appear to alter either the levels or subcellular localisation of EcR (D, F, H, J). Scale bars, 100 μm.

